# PRK introns provide modular tools for enhanced transgene expression in *Chlamydomonas reinhardtii*

**DOI:** 10.64898/2026.10.15.754060

**Authors:** Marta Bertolini, Lucile Jomat, Françoise Gilard, Bertrand Gakiere, Stéphane D. Lemaire, Pierre Crozet

**Affiliations:** Sorbonne Université, CNRS, Department of Computational, Quantitative and Synthetic Biology, F-75005 Paris, France; Sorbonne Université, CNRS, INSERM, Institut de Biologie Paris-Seine, F-75005 Paris, France; Sorbonne Université, Université de Technologie de Compiègne, CNRS, INSERM, Biofoundry Alliance Sorbonne Université, F-75005 Paris, France; Université Paris-Saclay, CNRS, INRAE, Université Evry, Institute of Plant Sciences Paris-Saclay, 91190, Gif sur Yvette, France; Université Paris Cité, CNRS, INRAE, Institute of Plant Sciences Paris-Saclay, 91190, Gif sur Yvette, France; Sorbonne Université, Polytech Sorbonne, F-75005 Paris, France

**Keywords:** Introns, PRK, *Chlamydomonas reinhardtii*, Synthetic biology, MoClo

## Abstract

*Chlamydomonas reinhardtii* is an attractive chassis for photosynthetic biotechnology, but reliable nuclear transgene expression remains a major limitation, despite continuing advances in genetic engineering. One effective strategy is the introduction of introns into transgene sequences to trigger Intron-Mediated Enhancement (IME). Genomic sequences retaining endogenous introns often show higher expression than their intronless counterparts, as we previously observed for the phosphoribulokinase (PRK) gene. Here, we investigated the IME activity of PRK introns, using the first intron of RBCS2 (RBCS2i1), the current reference IME element in Chlamydomonas, as a benchmark. The first two PRK introns, PRKi1 and PRKi2, emerged as the most active candidates. We assessed their IME activity using two reporter systems, two genetic backgrounds, and multiple promoter–terminator combinations, and observed enhancements up to 35-fold relative to their intronless counterparts. Finally, we validated their performance using patchoulol synthase, a challenging transgene whose expression requires introns and enables the production of the high-value sesquiterpene patchoulol. PRKi1 enabled detectable patchoulol production under conditions in which a single RBCS2i1 intron was insufficient. Overall, our results establish PRKi1 and PRKi2 as versatile, MoClo-compatible IME tools and provide a modular, Biofoundry-compatible framework for optimizing transgene expression for synthetic biology and biotechnological applications in *C. reinhardtii*.

## Introduction

*Chlamydomonas reinhardtii* is a photosynthetic unicellular eukaryotic green alga that serves as a well-established model organism for fundamental research, especially in the field of photosynthesis (Salomé and Merchant 2019).

Our laboratory has led the development of the Modular Cloning (MoClo) toolkit for *C. reinhardtii* (Crozet et al. 2018), which enables rapid and standardized Golden Gate assembly of transcriptional units from interchangeable genetic parts (Engler et al. 2014). This modular framework has greatly facilitated systematic testing and engineering of transgene expression in *C. reinhardtii*.

Despite these tools, expressing transgenes in *C. reinhardtii* remains challenging, particularly when high expression levels are required (Jaeger, Baier, and Lauersen 2019; Kyle J. Lauersen 2019; Rasala and Mayfield 2011). Several complementary strategies have been developed to address this limitation, including identification of stronger promoters (Schroda 2019; Schroda, Beck, and Vallon 2002), use of specific strains to increase transgene expression and/or reduce gene silencing (Kurniasih et al. 2016; Neupert, Karcher, and Bock 2009; Schroda 2019), codon usage optimization (Fuhrmann, Oertel, and Hegemann 1999; Schroda 2019) and introduction of introns (Baier et al. 2020; Eichler-Stahlberg et al. 2009; Lumbreras, Stevens, and Purton 1998). This last strategy exploits a phenomenon known as Intron-Mediated Enhancement (IME). IME designates any situation in which the presence of an intron increases transgene expression relative to an intronless counterpart (Gallegos and Rose 2015). This phenomenon has been widely documented across many eukaryotes, including yeast (Furger et al. 2002; Moabbi et al. 2012), nematodes (Okkema et al. 1993), insects (Bingham et al. 1988; Gasch, Hinz, and Renkawitz-Pohl 1989), mammals (Shaul 2017), land plants (Callis, Fromm, and Walbot 1987) and *C. reinhardtii* (Baier et al. 2020; Jaeger, Baier, and Lauersen 2019; Lumbreras, Stevens, and Purton 1998; Salomé and Merchant 2019). To activate IME, introns must be located within the transcribed region, and the effect is strongest when the intron is located close to the transcription start site (Callis, Fromm, and Walbot 1987; Clancy et al. 1994; Curi, Chan, and Gonzalez 2005; Laxa 2016; McElroy et al. 1990; Parra et al. 2011; Zalabák and Ikeda 2020). Although IME has been widely exploited to enhance transgene expression, its molecular basis remains poorly understood and appears highly dependent on the intron sequence and genetic context (Gallegos and Rose 2015; Laxa 2016). Recent work in *C. reinhardtii* has evaluated intron activity in standardized experimental frameworks, but only a limited subset of introns has been investigated (Xing et al. 2026). The most widely used IME elements in *C. reinhardtii* is the first intron of the RBCS2 gene (RBCS2i1), which confers a 5.5-fold increase in expression relative to an intronless control, when inserted as a single copy within a reporter gene (Baier et al. 2020). When present in multiple copies, an even greater enhancement can be achieved (Baier et al. 2018).

To facilitate the use of introns for transgene expression, a widely adopted bioinformatic tool called Intronserter (Jaeger, Baier, and Lauersen 2019) optimizes the number and positioning of introns, typically multiple copies of RBCS2i1, to achieve higher gene expression. Although powerful, simple and versatile, this strategy requires redesign of the coding sequence and often *de novo* synthesis, which is undesirable when rapid interchangeability of genetic parts is needed. An alternative and complementary strategy would therefore be to expand the repertoire of potent, standardized introns that can be rapidly interchanged without redesigning the coding sequence. Such standardized intron parts could facilitate rapid combinatorial testing, sharing between laboratories, Golden Gate shuffling and automated assembly in Biofoundry workflows (Crozet et al. 2018; Douchi et al. 2026). Previous studies have shown substantial variation in IME activity among *C. reinhardtii* introns, with some introns matching or exceeding RBCS2i1 in specific contexts (Baier et al. 2020). We previously showed that high-level expression of the *C. reinhardtii* Phosphoribulokinase (PRK) gene can be achieved with the genomic sequence but not with the corresponding intronless sequence (Boisset et al. 2023), strongly suggesting that PRK introns possess IME activity. In the present study, we therefore evaluated PRK introns (PRKi) as candidate IME elements and benchmarked their activity against RBCS2i1. Moreover, because IME activity is highly context-dependent, expanding the repertoire of standardized intron parts may provide alternative solutions for transgenes that perform poorly with IME elements currently available for genetic engineering.

To evaluate PRK introns as modular IME elements, we first benchmarked all PRK and RBCS2 introns against an intronless control using NanoLuc, a luciferase reporter whose high sensitivity and wide dynamic range allow enhancement to be quantified over several orders of magnitude (Crozet et al. 2018; Hall et al. 2012). We then characterized the leading candidates, PRKi1 and PRKi2, across different genetic backgrounds and regulatory contexts, using the yellow fluorescent protein mVenus as an alternative reporter and the challenging non-reporter transgene patchoulol synthase. Altogether, our data identify PRKi1 and PRKi2 as potent IME elements for transgene expression in *C. reinhardtii*, establish them as validated MoClo-compatible intron parts that expand the genetic engineering toolbox available to the community, and provide a MoClo- and Biofoundry-compatible workflow for the systematic screening and characterization of intron candidates for IME activity.

## Results

### PRKi1, PRKi2, and RBCS2i1 IME activity distribution in single clones

To investigate whether PRK introns contribute to the higher expression observed with the endogenous genomic PRK sequence (Boisset et al. 2023), the first two PRK introns were cloned as MoClo level 0 parts. These introns were selected because first introns have been reported to display the strongest IME activity in Arabidopsis (Gallegos and Rose 2015).

The introns were assembled upstream of the NanoLuc reporter. Each construct contained the intron followed by the NanoLuc coding sequence lacking its ATG, thereby requiring insertion of a linker sequence. This 24 nt linker (Figure 1a) contained an ATG and a G immediately upstream of the 5’ splice site, a configuration previously shown to support optimal IME activity (Baier et al. 2020). Two controls were also built: an intronless construct and a construct containing RBCS2i1, the reference IME intron in *C. reinhardtii*. All transcriptional units (TU) were under the control of the same regulatory elements P_PSAD_ and T_PSAD_ as promoter/5’UTR and 3’UTR/terminator, respectively (Figure 1b). To enable transformant selection, these constructs were combined with a TU conferring hygromycin resistance and transformed into the CC-5325 strain.

**Figure 1:**
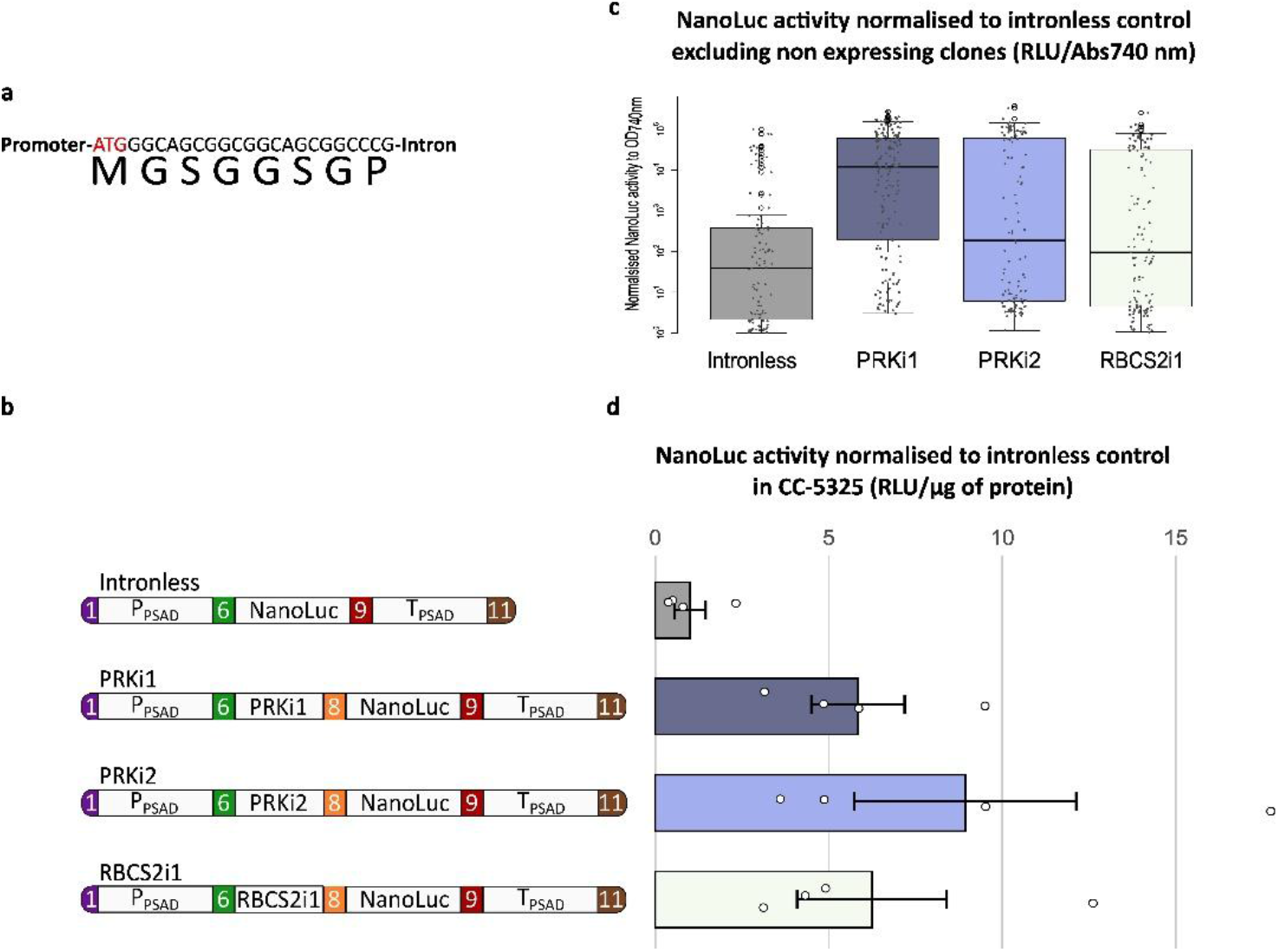
PRKi1 and PRKi2 mediate Intron-Mediated Enhancement of NanoLuc expression. (a) Sequence of the linker preceding all introns used. The start codon ATG is marked in red, and the G of the 5’ splice site is in green. (b) Schematic representation of the constructs used for NanoLuc transformation. P_PSAD_: PSAD promoter and 5’UTR; PRKi1: first intron of the phosphoribulokinase (PRK) gene; PRKi2: second intron of the PRK gene; RBCS2i1: first intron of gene 2 coding for the small subunit of Rubisco; NanoLuc: coding sequence of NanoLuc luciferase; T_PSAD_: PSAD 3’UTR and terminator. All introns are preceded by the linker sequence presented in panel (a). The numbers in colored boxes represent the MoClo fusion sites of level 0 (Crozet et al. 2018) (c) NanoLuc expression normalized by Abs_*740nm*_ of 300 independent clones from 3 independent transformations. (d) The bar plot represents NanoLuc activity for the different constructs. NanoLuc activity (RLU/µg protein) is normalized to the intronless control. Bars represent the mean ± SEM (N=4, biological replicates), all data are presented (white dots).

To assess the distribution of transgene expression, 300 colonies for each construct, derived from three independent transformations, were picked and NanoLuc activity was measured. Luminescence values were then normalized to chlorophyll autofluorescence to allow comparison among clones. The results presented in Figure 1c show that all tested introns enhanced NanoLuc expression. In particular, the frequency of clones expressing NanoLuc above a defined threshold (about 10-fold higher than the non-transformed control) was greater in the presence of each of the introns. While only 27% of the clones obtained with the intronless control exhibited NanoLuc activity above this threshold, this proportion rose to 96%, 84% and 83% for PRKi1, PRKi2 and RBCS2i1, respectively. Among the introns tested, PRKi1 yielded 5-fold more strongly expressing clones (defined as clones expressing more than 100-fold above the intronless control) than the intronless control, whereas PRKi2 and RBCS2i1 yielded 3-fold and 2.5-fold more such clones, respectively, with broader distributions and lower median expression values. Correct intron splicing was verified by RT-PCR on cDNA from three clones per construct and compared with PCR performed on genomic DNA, revealing the expected size difference on agarose gel (Figure S1). This shows that the introns are correctly spliced even in synthetic constructs and are functional as standalone parts.

### PRKi1 and PRKi2 enhance transgene expression in pooled transformants

To evaluate the impact of each intron on transgene expression independently of position effects arising from the genomic insertion site of the transgene, the same constructs (Figure 1b) were used to transform the CC-5325 strain (Lodha, Schulz-Raffelt, and Schroda 2008). All the colonies from each transformation plate were pooled, and proteins were then extracted and quantified prior to luminescence determination (Figure 1d).

NanoLuc measurements confirmed the trends observed in individual transformants (Figure 1c): PRKi1 and PRKi2 promoted IME to an extent comparable to or greater than RBCS2i1, resulting in 6-fold and 8-fold increases, respectively, compared to the intronless control. Importantly, the magnitude of enhancement observed for RBCS2i1 was consistent with previously published results, despite the use of a different readout system, here luminescence as opposed to antibiotic resistance in previous studies (Baier et al. 2018). These data demonstrate the IME activity of PRKi1 and PRKi2.

### Only the first introns of PRK and RBCS2 show strong IME activity

Because PRKi1 and PRKi2 displayed an IME activity comparable to RBCS2i1, we next tested the five remaining PRK and the two additional RBCS2 introns. The constructs were built using the same general design as above, with the corresponding introns combined with the NanoLuc reporter construct (Figure 2a). After transformation of CC-5325 cells, global NanoLuc activity was measured in pooled transformants (Figure 2b). These data showed that the first introns of PRK and RBCS2 (Figure 1) displayed the highest IME activity of all the introns tested. Accordingly, subsequent experiments focused on PRKi1, PRKi2, and RBCS2i1. What must be kept in mind is that the constructs of Figure 2 were transformed and assayed in an experimental series distinct from that of Figure 1. Although both series included their own intronless control and all values are expressed relative to that internal control, the absolute values of the two figures are not directly comparable.

**Figure 2:**
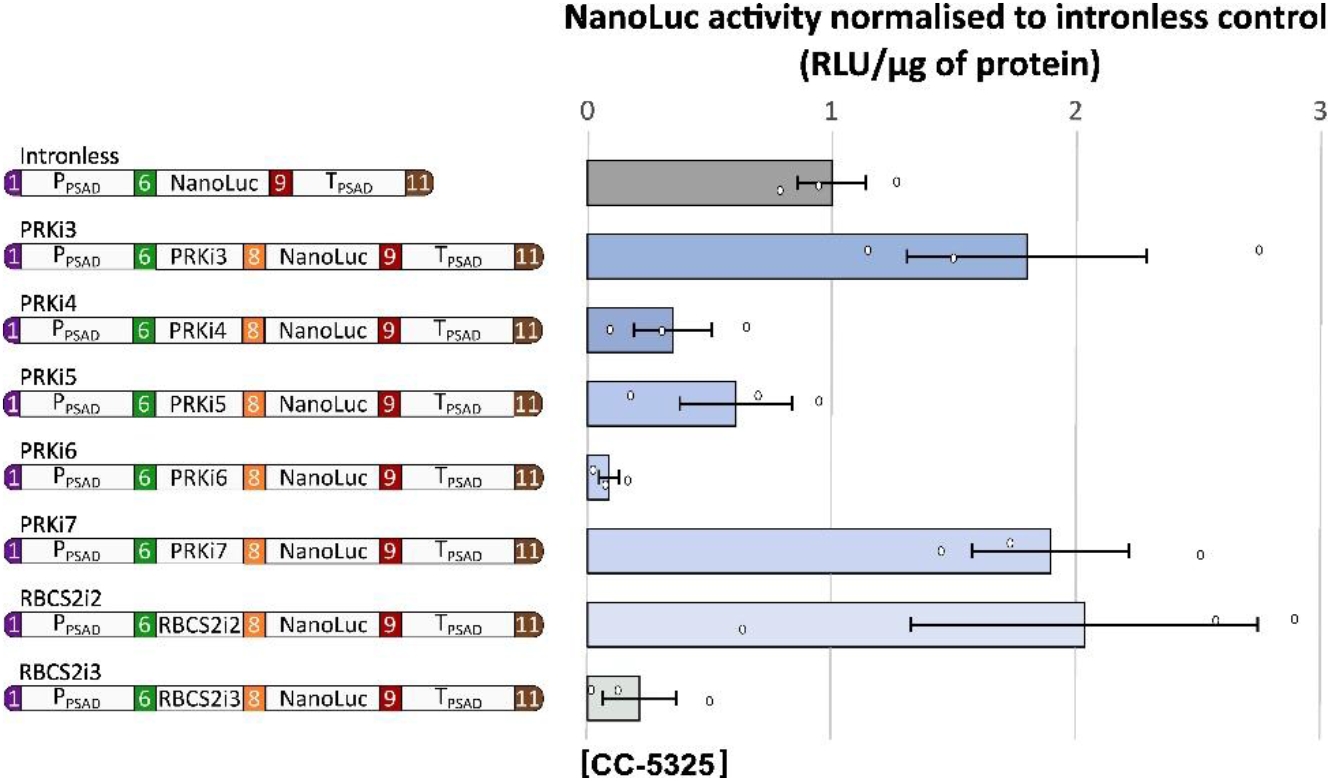
Evaluation of IME activity of additional PRK and RBCS2 introns. (a) Schematic representation of the constructs used for NanoLuc transformation with the other PRK and RBCS2 introns. P_PSAD_: PSAD promoter and 5’UTR; NanoLuc: coding sequence of NanoLuc luciferase; PRKi3: third intron of the phosphoribulokinase (PRK) gene; PRKi4: fourth intron of the PRK gene; PRKi5: fifth intron of the PRK gene; PRKi6: sixth intron of the PRK gene; PRKi7: seventh intron of the PRK gene; RBCS2i2: second intron of gene 2 coding for small subunit of Rubisco; RBCS2i3: third intron of gene 2 coding for the small subunit of Rubisco; T_PSAD_: PSAD 3’UTR and terminator. All introns are preceded by the linker sequence presented in Figure 1a. (b) NanoLuc activity (RLU/µg protein) of each intron-containing construct normalized to the intronless control. Bars represent the mean ± SEM (N=3, biological replicates), all data are presented (white dots).

### IME activity depends on promoter and terminator context

To further evaluate the context dependence of PRKi1 and PRKi2 IME activity, we tested these introns in alternative regulatory environments. For this purpose, the three selected introns were assembled with different promoters and terminators (Figure 3a and c). These constructs retained the same overall architecture as the original NanoLuc construct (Figure 1b), with only the promoter or terminator being changed. In both series, the P_PSAD_ promoter was replaced by the P_AR_ promoter, a chimeric promoter derived from HSP70A and RBCS2 promoters (Schroda, Beck, and Vallon 2002), which allows strong expression in *C. reinhardtii* (Schroda, Blöcker, and Beck 2000). In the second series, the T_PSAD_ was replaced by the RPL23 terminator, which contains an intron in its 3’UTR and was recently reported to support improved transgene expression (Aschern et al. 2025). This 3’UTR intron could contribute to IME independently of the intron placed upstream of the coding sequence. However, T_RPL23i_ is present in all four constructs of this series, including the intronless control, so that the relative comparison within the series remains internally valid. As above, *C. reinhardtii* cells were transformed with these constructs, transformants were pooled to reduce position effects, and NanoLuc activity was measured in total soluble protein extracts (Lodha, Schulz-Raffelt, and Schroda 2008).

**Figure 3:**
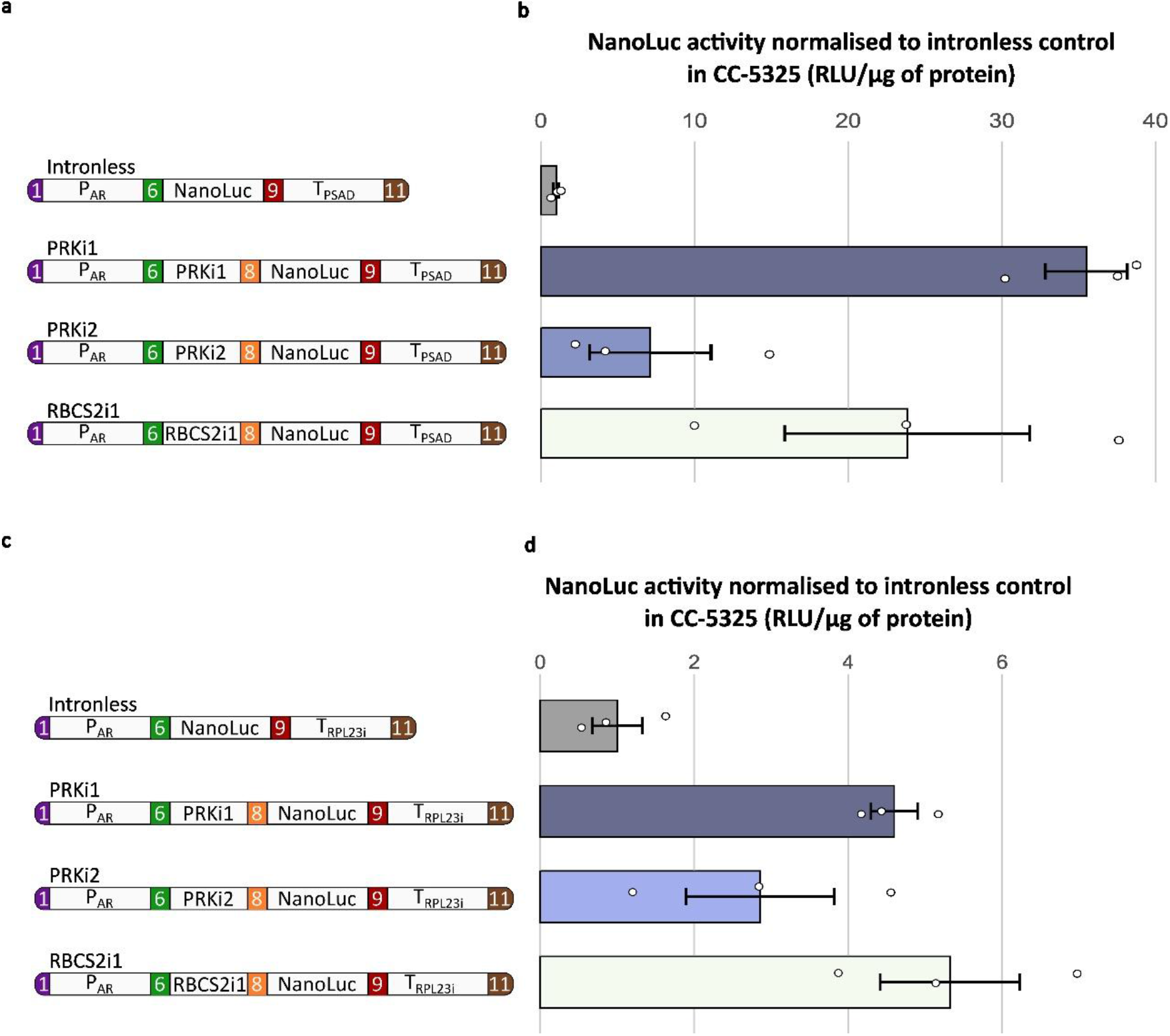
IME activity depends on promoter and terminator context. (a-c) Schematic representation of the constructs used for NanoLuc transformation with different promoter and terminator combinations. P_AR_: synthetic promoter and 5’UTR generated by fusion of the HSP70A promoter and the RBCS2 promoter; PRKi1: first intron of the phosphoribulokinase (PRK) gene; PRKi2: second intron of the PRK gene; RBCS2i1: First intron of gene 2 coding for the small subunit of Rubisco; NanoLuc: coding sequence of NanoLuc luciferase; T_PSAD_: PSAD 3’UTR and terminator; T_RPL23i_: RPL23 terminator and 3’UTR including one intron. The 3 tested introns are preceded by the linker sequence presented in Figure 1a. (b-d) NanoLuc activity (RLU/µg protein) for the different intron-containing constructs normalized to the intronless control. Bars represent the mean ± SEM (N=3, biological replicates), all data are presented (white dots).

Changing the promoter/5’UTR to P_AR_ markedly affected the extent of IME (Figure 3b), with PRKi1 yielding a 35-fold increase compared to the corresponding intronless control. By contrast, substitution of the 3’UTR/terminator yielded a similar, although slightly lower, range of IME values than those observed with T_PSAD_ (Figure 1d, 3d).

### PRKi1 and PRKi2 remain active in a different strain

Because IME is highly context-dependent and multiple *C. reinhardtii* strains are used across the field, we next examined PRKi1 and PRKi2 IME activity in an alternative genetic background. The SRTA-2 strain was selected because it carries a knockout of the gene encoding a putative Sir2-type histone deacetylase, which attenuates transgene silencing (Einhaus et al. 2025; Li et al. 2019; Neupert et al. 2020). This background is increasingly used in the Chlamydomonas community, making it a relevant context in which to assess the performance of PRKi1 and PRKi2.

After transformation of SRTA-2 with the constructs shown in Figure 4a, colonies from three independent transformations were pooled to prevent position effects, yielding more than 300 clones in total, prior to protein extraction (Lodha, Schulz-Raffelt, and Schroda 2008). Although the overall pattern of IME activity was similar to that observed in CC-5325, the fold-increase relative to the intronless control was lower in the SRTA-2 background (Figure 4b).

**Figure 4:**
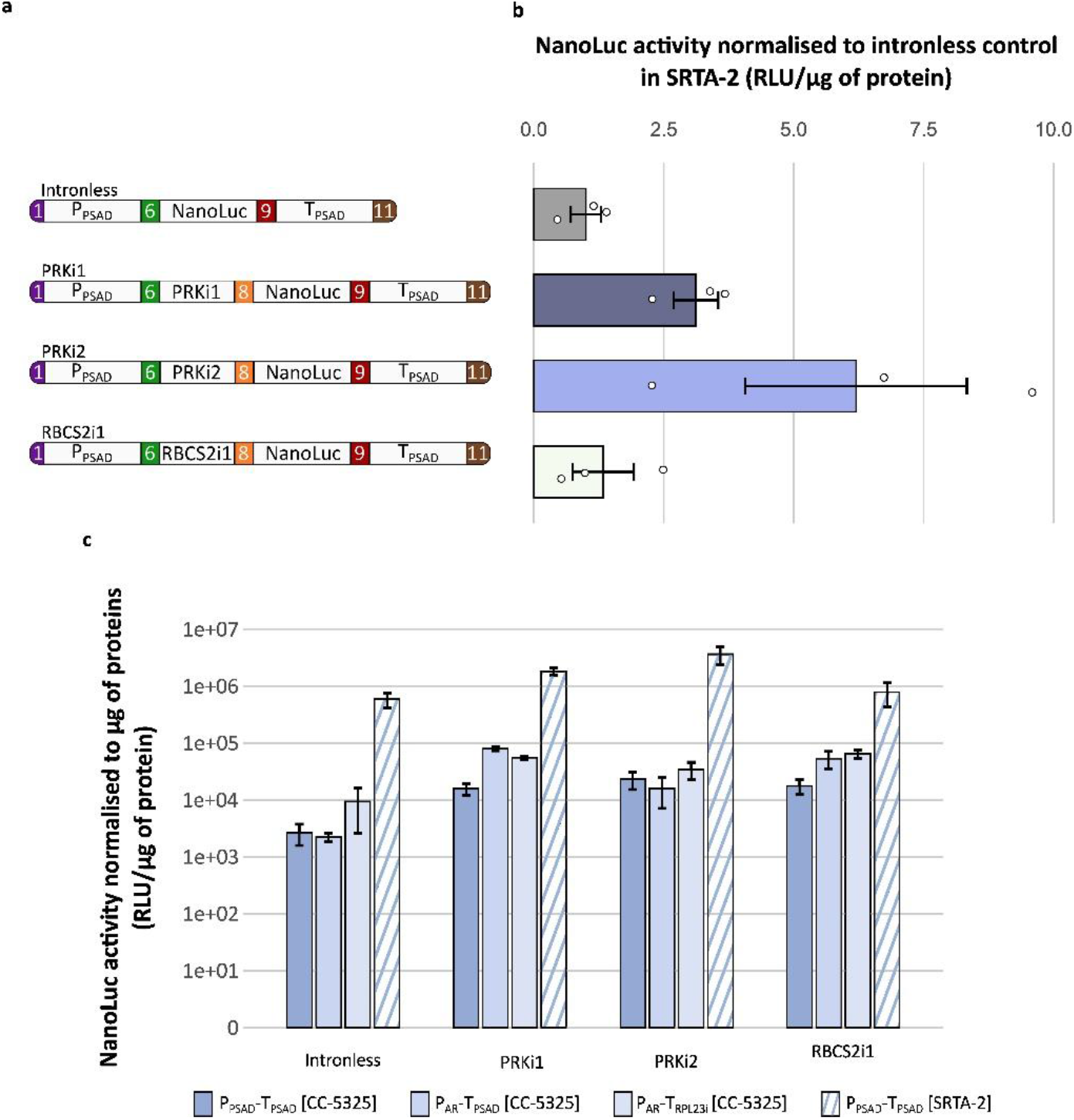
PRKi1 and PRKi2 remain active in the SRTA-2 strain. (a) Schematic representation of the constructs used for NanoLuc transformation. P_PSAD_: PSAD 5’UTR promoter; PRKi1: first intron of the phosphoribulokinase (PRK) gene; PRKi2: second intron of the PRK gene; RBCS2i1: First intron of gene 2 coding for the small subunit of Rubisco; NanoLuc: coding sequence of NanoLuc luciferase; T_PSAD_: PSAD 3’UTR and terminator. All introns are preceded by the linker sequence presented in Figure 1a. (b) The bar plot represents NanoLuc activity (RLU/µg protein) for the different constructs, normalized to the intronless control. Bars represent the mean ± SEM (N=3, biological replicates), all data are presented (white dots). (c) Summary of NanoLuc results obtained by testing different constructs (Figures 2, 3 and 4a) and different strains.

We then compared the full NanoLuc datasets to identify the combinations yielding the highest expression in Chlamydomonas (Figure 4c). As expected, the SRTA-2 strain provided a highly permissive background for transgene expression. Adding PRKi1 or PRKi2 to a transcriptional unit containing P_PSAD_ and T_PSAD_ further increased expression, even in this strain, and the highest expression was observed with PRKi2 with a 6-fold improvement. In the CC-5325 background, the highest NanoLuc expression was obtained with PRKi1 in combination with P_AR_ and T_PSAD_, resulting in a 35-fold increase compared to the intronless counterpart.

### IME activity is lower with mVenus than with NanoLuc

Because fluorescent proteins are widely used to monitor transgene expression and subcellular localization, we next tested whether PRKi1 and PRKi2 could enhance expression of mVenus. Previous work showed that tagging the PcPS protein with the mVenus yellow fluorescent protein can increase transgene expression when introns are present (Baier et al. 2020), prompting us to ask whether mVenus expression would also benefit from PRKi1- and PRKi2-mediated IME. To this end, constructs analogous to the NanoLuc reporter constructs were generated, with the NanoLuc coding sequence replaced by mVenus coding sequence without any intron (Figure 5a) and introduced into CC-5325. As in previous experiments, colonies from three independent transformations were pooled, and proteins were extracted and quantified. IME activity was markedly lower with mVenus than with NanoLuc with a maximum enhancement of 2.5-fold (Figure 5b). This confirms that IME is strongly context-dependent, including dependence on the coding sequence being expressed.

**Figure 5:**
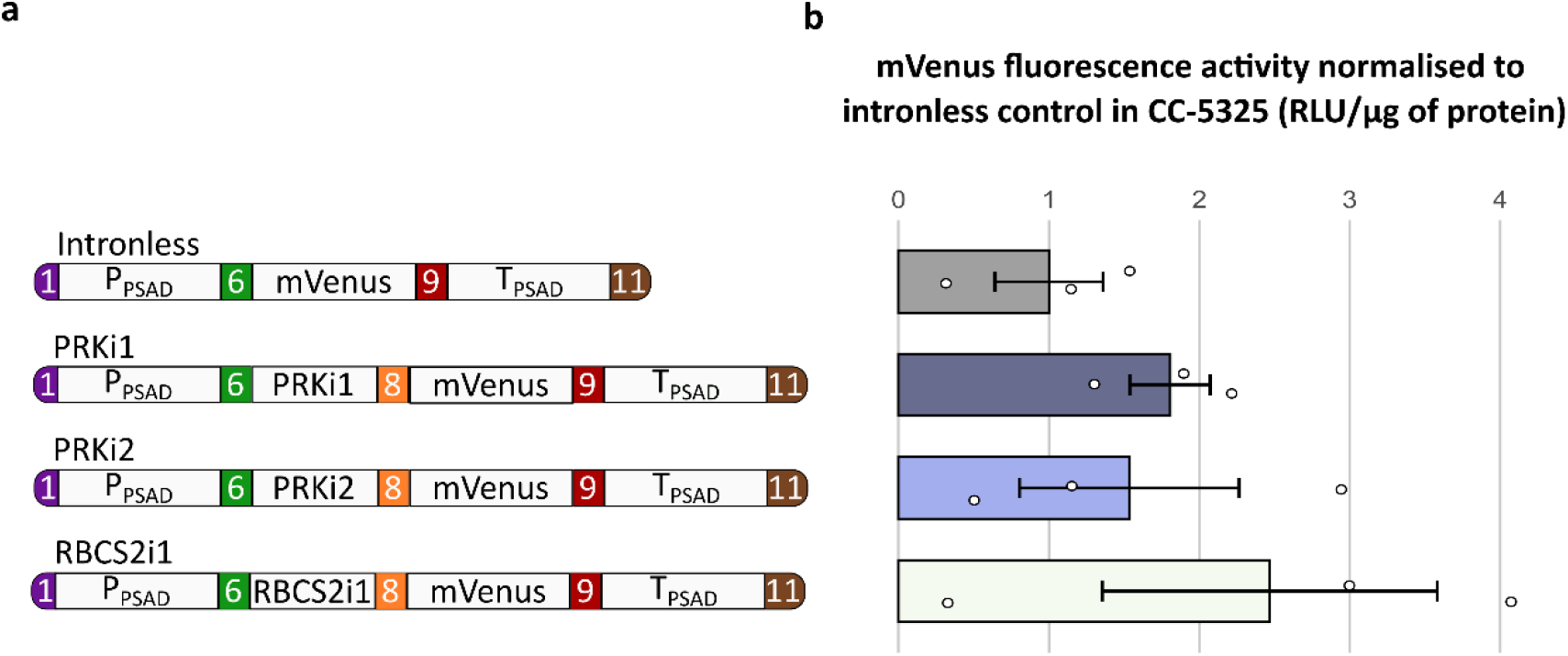
Evaluation of IME activity of PRKi1, PRKi2, and RBCS2i1 on mVenus expression. (a) Schematic representation of the constructs used for mVenus transformations. P_PSAD_: PSAD promoter and 5’UTR; RBCS2i1: first intron of the RBCS2 gene; PRKi1: first intron of the PRK gene; PRKi2: second intron of the PRK gene; T_PSAD_: PSAD 3’UTR and terminator. (b) The bar plot represents mVenus fluorescence (RFU/µg protein) for the different constructs, normalized to the intronless control. Bars represent the mean ± SEM (N=3, biological replicates), all data are presented (white dots).

### PRKi1 supports patchoulol synthase expression and patchoulol production

Finally, to validate the activity of PRKi1 and PRKi2 in the context of a non-reporter transgene, we tested their ability to enhance expression of patchoulol synthase (PcPS). PcPS expression enables the production, in *C. reinhardtii*, of patchoulol, a valuable sesquiterpene that can be readily quantified by GC-MS. Because this gene is difficult to express in the absence of introns, it has previously been used as a stringent test of IME activity in *C. reinhardtii* (Baier et al. 2020). We therefore evaluated the two PRK introns in this demanding expression context using four constructs (Figure 6a). To maintain a simple and modular design, without requiring re-synthesis of the sequence to incorporate introns, a single intron was inserted upstream of the coding region. Because larger reporter tags such as mVenus increase transgene complexity, we instead used a HiBiT tag. HiBiT is an 11-amino-acid peptide tag derived from split NanoLuc that generates bioluminescence upon addition of the rest of the enzyme (LgBiT) and substrate in solution (Inckemann et al. 2025). The HiBiT tag enabled rapid and quantitative screening of transformants prior to GC-MS analysis of patchoulol production.

**Figure 6:**
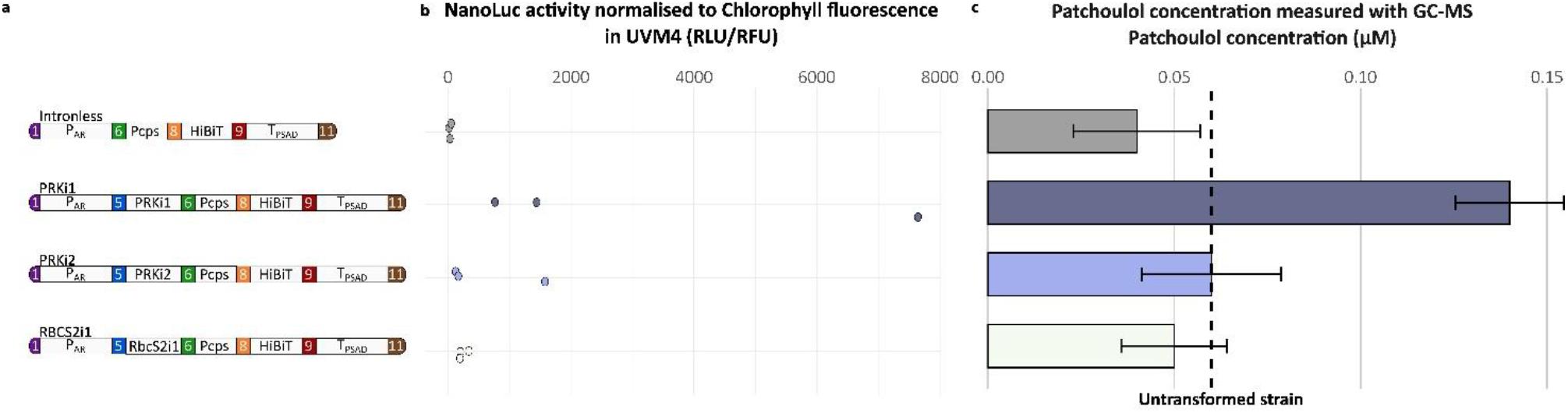
Results of patchoulol production after GC-MS analysis. (a) Schematic representation of the constructs used for PcPS transformations. P_PSAD_: PSAD promoter and 5’UTR; RBCS2i1: first intron of the RBCS2 gene; PRKi1: first intron of the PRK gene; PRKi2: second intron of the PRK gene; T_PSAD_: PSAD 3’UTR and terminator. (b) Representation of the NanoLuc activity in UVM4 measured using the HiBiT tag (RLU) normalized to chlorophyll autofluorescence (RFU). The data shown are for the three best clones that were then tested for patchoulol production. (c) The bar plot represents the average concentration of patchoulol in three clones previously identified based on their HiBiT signal. Bars represent the mean ± SEM (N=3, biological replicates).

The four constructs were introduced into the UVM4 strain, the same background used previously by T. Baier and coworkers (Baier et al. 2020). For each construct, at least three clones showing significant HiBiT signal were identified (Figure 6b), and the three best clones per construct were grown in flasks with a dodecane overlay. Patchoulol production was then quantified by GC-MS analysis (Figure 6c). As expected, neither the intronless construct nor the construct containing a single copy of RBCS2i1 supported detectable expression of PcPS. Likewise, a single copy of PRKi2 did not support substantial PcPS expression, although patchoulol could be detected and quantified, at levels twice those of the intronless control (Figure 6c). In contrast, a single copy of PRKi1 supported substantial patchoulol production (Figure 6c). These results are consistent with previous reports showing that a single copy of RBCS2i1 is insufficient to support detectable patchoulol production (Baier et al. 2020), and suggest that PRKi1 may confer stronger IME activity in this specific context, since it was the only intron tested for which patchoulol could be reproducibly detected and quantified. We emphasize that this conclusion is qualitative as it rests on three clones per construct selected on the basis of their HiBiT signal, and the difference between constructs was not tested statistically.

## Discussion

In this study, we validated PRKi1 and PRKi2 as modular intron parts that mediate robust IME for nuclear transgenes in *Chlamydomonas reinhardtii*. The mechanism of IME remains incompletely understood in every organism in which it has been described (Gallegos and Rose 2015; Laxa 2016), and the practical value of well-characterized intron parts does not depend on that mechanism being resolved. We achieved this by systematically testing genetic circuits that combined two reporter genes, a non-reporter gene that is particularly difficult to express, and several regulatory elements, in three *C. reinhardtii* strains. Moreover, we validated NanoLuc as a reporter for IME studies and demonstrated the *in vivo* use of the split NanoLuc HiBiT-LgBiT system expressed from the nuclear genome.

Although the improvement in expression mediated by PRKi1 and PRKi2 was significant in all tests performed, the magnitude of the enhancement varied markedly between constructs and conditions. This variability is consistent with previous reports showing that IME depends on the sequence position of the intron, and on the surrounding promoter, terminator and coding sequence (Baier et al. 2020; Laxa 2016). This underlines the value of expanding the intron toolbox so that IME can be exploited across a broad range of contexts to improve transgene expression.

Much of the spread observed in our datasets is inherent to nuclear transformation in *C. reinhardtii*. Random integration generates position effects, variable copy number and variable susceptibility to silencing, so that expression levels among individual transformants of a single construct routinely span two to three orders of magnitude (Figure 1c). We addressed this in two complementary ways: by analyzing the full distribution of 300 independent clones per construct, which describes this variability rather than averaging it, and by analyzing pooled transformants, which averages it at the cost of masking clone-to-clone differences. The residual variability between biological replicates of pooled transformants reflects differences in transformation efficiency and in the number of clones recovered per plate, both of which alter the composition of the pool. To limit these effects, all constructs of a series were transformed and processed in parallel, and every value is expressed relative to the intronless control of the same series.

Accordingly, analysis of NanoLuc expression in pooled transformants (Figure 4c) allows us to draw several general conclusions. The highest absolute NanoLuc expression was obtained in the SRTA-2 strain, particularly with PRKi2 in combination with the PSAD promoter and terminator, in line with the reduced transgene silencing reported for this background (Einhaus et al. 2025; Neupert et al. 2020). However, in the widely used CC-5325 strain, the best performing configuration combined PRKi1 with the AR promoter and PSAD terminator yielding a 35-fold increase relative to the corresponding intronless construct, and performing comparably or better than RBCS2i1 coupled to AR promoter and either the RPL23 or PSAD terminator. These observations highlight that PRKi1 and PRKi2 can match or even surpass the performance of RBCS2i1 in appropriate regulatory contexts. Figure 4c also shows that the genetic background (here CC-5325 and SRTA-2) exerts a larger effect on absolute transgene expression than the choice of intron. IME and silencing suppression are therefore complementary rather than interchangeable strategies, and comparisons between introns are only meaningful within a similar genetic background. This also implies that the fold-enhancement values reported here, like all IME values in the literature, are conditional on the strain in which they were measured.

The patchoulol synthase experiments further underscore this context specificity and demonstrate that PRKi1 can function as a practical IME tool for challenging, non-reporter genes. As previously reported, a single copy of RBCS2i1 is insufficient to support detectable patchoulol production (Baier et al. 2020), and this finding was reproduced here: neither the intronless PcPS construct nor the PcPS construct containing a single RBCS2i1 intron yielded measurable patchoulol. Similarly, a single PRKi2 intron did not produce substantial PcPS activity. In contrast, PcPS constructs containing a single PRKi1 intron supported detectable patchoulol production, indicating that PRKi1 can mediate effective IME even in a stringent biotechnological context that is highly sensitive to intron choice. The absolute patchoulol titers obtained here remain modest compared with those reported for strains engineered specifically for sesquiterpenoid production, which is expected. Our constructs carry a single intron rather than a fully intron-optimized synthetic coding sequence, they were not combined with any metabolic engineering of the isoprenoid precursor supply, and the cultures were grown for six days at 5 mL scale in multi-well plates rather than in optimized photobioreactors. The purpose of this experiment was to test whether a single modular intron is sufficient to render a notoriously difficult transgene functional, not to maximize product titer.

Our data also illustrate that the extent of IME is strongly influenced by the regulatory environment flanking the intron. Substitution of the PSAD promoter with the chimeric AR promoter had a pronounced impact on IME, particularly for PRKi1, leading to markedly higher NanoLuc expression than PSAD-based constructs. Conversely, replacement of the PSAD terminator with the RPL23 terminator, which itself contains a 3′ UTR intron (Aschern et al. 2025), maintained IME within a similar range, suggesting that promoter choice is a dominant determinant of IME amplitude in our system, whereas the tested 3′ UTR/terminator variants exerted more modest effects.

The comparison of NanoLuc and mVenus reporters confirmed that IME strength is not an intrinsic, reporter-independent property of a given intron but rather depends on the coding sequence and possibly on mRNA features specific to the reporter. For mVenus, IME mediated by PRKi1, PRKi2, or RBCS2i1 was substantially weaker than for NanoLuc under otherwise similar conditions, supporting the view that IME results from an interplay between intron and transcript architecture rather than a simple additive “boost” independent of context (Baier et al. 2020; Laxa 2016).

On the basis of the whole dataset, we can offer the following practical guidance. In the widely used CC-5325 background, PRKi1 combined with the AR promoter and the PSAD terminator gave the largest enhancement and is our recommended starting point. In silencing-suppressed backgrounds such as SRTA-2, PRKi2 combined with PSAD regulatory elements performed best. RBCS2i1 remains a sound default and was not consistently outperformed across the conditions tested. The value of PRKi1 and PRKi2 is that they provide alternatives in the contexts where RBCS2i1 is insufficient, as illustrated by the patchoulol synthase constructs. For a difficult transgene, we recommend testing PRKi1, PRKi2 and RBCS2i1 in parallel rather than choosing one intron in advance, a comparison that the modular design described here reduces to a single round of Golden Gate assembly.

Several of the introns tested here reduced rather than increased expression relative to the intronless control (Figure 2). Negative IME has been reported before in *C. reinhardtii*, notably for βTUB2i1, which enhanced expression in the original MoClo study (Crozet et al. 2018) but was found to be detrimental by Baier *et al*. (2020). Since βTUB2i1 is the intron currently distributed in the MoClo kit, this discrepancy reinforces the point that an intron cannot be assumed to be beneficial outside the context in which it was characterized, and argues for systematically including an intronless control when a new transcriptional unit is designed.

From a synthetic biology perspective, a key advantage of PRKi1 and PRKi2 is their compatibility with modular cloning strategies. In our designs, a single intron is inserted immediately upstream of the coding sequence *via* standardized MoClo parts, avoiding the need to re-synthesize or re-order the coding sequence, in contrast to approaches that distribute multiple introns throughout the coding sequence (Jaeger, Baier, and Lauersen 2019). This modularity facilitates combinatorial testing of introns, promoters, and terminators within Biofoundry workflows and enables sharing of characterized intron parts across the community (Douchi et al. 2026). Our results indicate that PRKi1, in particular, is a strong candidate for inclusion as a standard IME element, alongside RBCS2i1, in future Chlamydomonas toolkits. This approach complements rather than replaces Intronserter and RBCS2i1. Intronserter remains the method of choice when a transgene is synthesized *de novo* and maximal expression is the priority, since distributing multiple introns throughout a coding sequence can deliver enhancements beyond those achievable with a single upstream intron. RBCS2i1 remains the best-characterized and most widely validated IME element in *C. reinhardtii*. The single-part strategy described here is most useful when an existing coding sequence must be kept as it is, when several introns must be compared quickly, or when an intron is needed as an interchangeable module in a combinatorial assembly. Consistent with this, we propose that PRKi1 and PRKi2 be deposited at the Chlamydomonas Resource Center for inclusion in the existing MoClo kit rather than distributed as a separate kit, so that users are not required to purchase an additional plate to access them.

In practice, the combinations identified here can be further expanded through MoClo-based part shuffling and systematically screened at high throughput using Biofoundry infrastructures to identify optimal intron– promoter–terminator sets for specific applications.

In conclusion, we establish PRKi1 and PRKi2 as effective and modular IME elements for nuclear transgene expression in *Chlamydomonas reinhardtii*. Their compatibility with standardized MoClo assembly enables rapid combinatorial optimization of transgene expression without redesigning the coding sequence. Together with the successful expression of the challenging patchoulol synthase transgene, these results expand the synthetic biology toolbox available for engineering *C. reinhardtii* and provide a Biofoundry-compatible framework for the systematic identification of expression configurations tailored to specific biotechnological applications.

## Experimental procedures

Unless otherwise indicated, all reagents were purchased from Merck (St. Louis, MO, USA).

### Chlamydomonas reinhardtii strains, medium and culture conditions

Three *Chlamydomonas reinhardtii* strains were used in this study. CC-5325 is the reference strain used for transformation with the NanoLuc and mVenus constructs. This strain is also known as CMJ030 and is the background for all the clones coming from the first genome-wide mutant library of *Chlamydomonas reinhardtii* (CLiP) (Blaby et al. 2014; Li et al. 2019; Zhang et al. 2014). SRTA-2 (Li et al. 2019; Neupert et al. 2020) was used as an additional genetic background to validate IME activity with the NanoLuc constructs. This strain is a CRISPR/Cas9-derived knockout of the gene encoding a putative Sir2-type histone deacetylase (SRTA), generated to reduce epigenetic transgene silencing and thereby improve transgene expression. It should not be confused with UVM4, which is a UV-mutagenesis-derived strain carrying the *srta-2* allele affecting the same silencing pathway (Neupert, Karcher, and Bock 2009). UVM4 was used for transformation with the PcPS constructs. All three strains are photosynthetically competent. CC-5325 was used as the reference background for transgene expression analysis, because it has not been selected or engineered for enhanced transgene expression, whereas SRTA-2 and UVM4 have. All results are interpreted in the light of these strain properties.

Strains were grown at 25°C in Tris-Acetate-Phosphate (TAP) (Gorman and Levine 1965). When antibiotic selection was required, the medium was supplemented with hygromycin at a final concentration of 20 μg/mL. Strains were maintained on solid TAP plates containing 1.5% agar as stock cultures, whereas liquid cultures were used for transformation and downstream analyses. Liquid cultures were maintained under agitation (120 rpm) in Erlenmeyer flasks under continuous white light (40-60 μmol photons m^−2^s^−1^).

### Plasmid construction

All constructs were validated *in silico* using Serial Cloner 2.6.1. Following sequence validation, plasmids were assembled according to the MoClo protocol (Crozet et al. 2018) for all three levels: p0, p1 and pM. The only modification to the published protocol was the replacement of 2 μL of New England Biolabs CutSmart Buffer plus 2 μL of 10 mM ATP (buffered with 100 mM Tris-HCl pH 7.9) by 2 μL of New England Biolabs T4 DNA ligase buffer and 2 μL H_2_O. The MoClo reaction was conducted in a thermocycler as previously reported (Crozet et al. 2018). When positive clones were difficult to obtain, the number of 37°C-16°C cycles was increased from three to five. All sequences used are listed in Supplementary Table S1. Characteristics and sequence files of the intron-containing level 0 plasmids, not included in the published MoClo toolkit (Crozet et al. 2018) are provided in Data S1 and S2.

### Escherichia coli strains and handling

*E. coli* NEB10β strain (NEB reference C3019H, New England Biolabs Ipswich, MA, USA) was used for cloning and plasmid amplification. Bacterial transformation was performed by heat-shock according to the manufacturer’s instructions. Briefly, 4 μL of the MoClo reaction were added to 20 μL of competent bacteria in a 1.5 mL microcentrifuge tube. The mixture was then incubated at 42°C for 45 s and then placed on ice for 4 minutes. Subsequently, 500 μL of LB medium were added, and bacteria were allowed to recover at 37°C with shaking at 500 rpm for 30 minutes. After recovery, bacteria were plated on selective LB-agar and incubated overnight at 37°C, or for 48 h at 25°C. Bacteria were grown in liquid or solid LB medium prepared from LB powder (Sigma-Aldrich: 102654303). According to the MoClo protocol (Crozet et al. 2018), ampicillin (100 μg/mL), kanamycin (50 μg/mL) or spectinomycin (50 μg/mL) was added to the media as appropriate for plasmid selection. For blue-white screening, X-gal was added at 80 μg/mL.

### Plasmid extraction, validation and digestion

Colonies growing on selective plates were inoculated into 5 mL LB containing the appropriate antibiotic, in 15 mL culture tubes. Cultures were grown overnight at 37°C or for 48h at 25°C to allow plasmid amplification. Cultures were centrifuged at 4000 *g* for 5 min and the plasmids were purified from the cell pellet using the Macherey-Nagel NucleoSpin® Plasmid Kit according to the manufacturer’s instructions. Purified plasmids were validated by restriction digestion and/or sequencing (Sanger or Nanopore, Eurofins genomics, France). For *C. reinhardtii* transformation, 1 µg plasmid DNA was digested with BsaI for 4h at 37°C. For protocol 1, the DNA was resolved by agarose gel electrophoresis and the insert was purified from the gel with the Macherey-Nagel PCR Cleanup Kit according to the manufacturer’s recommendations. For protocol 2, the DNA was heated at 85°C to denature the restriction enzyme and purified with the Macherey-Nagel PCR Cleanup Kit according to the manufacturer’s recommendations. All enzymes and buffers were purchased from New England Biolabs (Ipswich, USA).

### Chlamydomonas reinhardtii transformation

*Chlamydomonas reinhardtii* cells were transformed by electroporation using two distinct protocols, depending on the strain. In all cases, cells were harvested during exponential growth phase (1 to 5 million cells/mL) by centrifugation, resuspended in electroporation buffer, electroporated, and allowed to recover overnight (16 h) on a rotating wheel in 10 mL of recovery medium. After recovery, cells were harvested again and plated on selective solid medium. Protocol-specific parameters are reported in Table 1.

**Table 1.** Electroporation parameters used for transformation of the different *Chlamydomonas reinhardtii* strains.

| Protocol<br>(Strain) | Cells per transformation<br>(Initial cell concentration<br>at harvest and<br>centrifugation<br>parameters) | Electroporation<br>and recovery<br>medium | Electroporation<br>parameters | Electroporator | Cuvette<br>gap<br>size | DNA<br>used |
| --- | --- | --- | --- | --- | --- | --- |
| **Protocol 1**<br>(Crozet et<br>al. 2018)<br>**(CC-5325)** | 4.8x10<sup>7</sup> cells<br>(1-5x10<sup>6</sup> cells/mL<br>1500 g, 2 min) | TAP+ Sucrose 60<br>mM | 800 V, 25 µF | Bio-Rad Gene<br>pulser II | 0.4 cm | 200 ng<br>of<br>purified<br>insert |
| **Protocol 2**<br>(Yamano,<br>Iguchi, and<br>Fukuzawa<br>2013)<br>**(SRTA-2**<br>**and UVM4)** | 2.6x10<sup>7</sup> cells<br>(4-8x10<sup>6</sup> cells/mL<br>4000 g 3 min) | TAP + Sucrose<br>40 mM | 2x 200 V, 8 ms<br>pulses (50ms<br>interval)<br>5x 20 V, 50 ms<br>(50ms interval) | NEPAGENE 21 | 0.2 cm | 500 ng<br>of<br>digested<br>DNA |

All data reported below derive from at least three independent transformations. Within a given experimental series, all constructs (including the intronless control) were cloned in parallel, transformed on the same day, from the same batch of cells, with the same transformation mix and the same mass of DNA (Table 1), and processed together through selection, protein extraction and measurement. All comparisons reported in this study are therefore internal to a single series and are expressed relative to the intronless control of that same series. Absolute values should not be compared between figure panels corresponding to different series. Because intron-containing constructs are longer than their intronless counterparts, the use of equal DNA masses corresponds to a marginally lower molarity for intron-containing constructs (at most 5% for the constructs used here).

### Protein extraction and quantification

Cells were resuspended in 500 µL of ice-cold protein extraction buffer (30 mM Tris-HCl pH 7.9, 1 mM EDTA, Antiprotease complete tablets (1 tablet for 50 mL, Roche, Basel, CH)). Cells were lysed by four cycles of flash-freezing in liquid nitrogen followed by thawing at room temperature in a water bath. The cell debris were pelleted by centrifugation at 4°C at *21,000 g* for 20 min and the supernatant was collected. Protein concentration was determined using a bicinchoninic acid assay with Bovine Serum Albumin as the standard.

### NanoLuc and HiBiT assay

NanoLuc activity was measured using the Nano-Glo™ Luciferase Assay system (Promega, Madison, WI, USA). The reagent was prepared according to the manufacturer’s instructions by mixing 1 part of substrate with 50 parts of Nano-Glo buffer. Then, 30 μL of the mix were added to 30 μL of quantified protein extract. Measurements were performed in duplicate in white 96-well plates, and luminescence was recorded using a BMG Labtech CLARIOstar plate reader (Ortenberg, Germany) at 470 ± 20 nm.

The HiBiT assay was performed using the Nano-Glo® HiBiT Lytic Detection System (Promega, Madison, WI, USA) according to the manufacturer’s instructions. A reaction mix was prepared by combining 1 volume of LgBiT Protein, 2 volumes of Nano-Glo® HiBiT Lytic substrate and 100 volumes of Nano-Glo® HiBiT Lytic Buffer. Both assays were performed in duplicate in white 96-well plates. For NanoLuc, 1 µg, 5 µg and 10 µg of total crude protein extract were tested. For HiBiT, 30 μL of the mix were added to 30 μL of liquid culture and mixed by pipetting to promote cell lysis. Samples were then incubated for 10 minutes at room temperature on an orbital shaker at 300 rpm.

For data analysis, only measurements falling within the linear range of both the NanoLuc assay and the plate reader were retained. Accordingly, only relative luminescence units (RLU) between 10^3^ and 10^6^ were used for the analyses.

### mVenus fluorescence analyses

mVenus fluorescence was measured on protein extracts in black 96-well plates using a BMG Labtech CLARIOstar plate reader (Ortenberg, Germany). Excitation was set to 497 ± 15 nm and emission was recorded at 540 ± 20 nm. Fluorescence values were normalized to total protein content.

### Patchoulol production and quantification

Transformants were inoculated in 5 mL cultures in 6-well plates at a starting concentration of 10^6^ cells/mL and grown for 6 days under a 500 µl dodecane overlay. The dodecane phase was then harvested by centrifugation and a 50 µl aliquot was supplemented with 50 μL of sclareol at 20 µmol/ml as an internal standard prior to GC/MS analysis.

One μL of sample was injected in splitless mode on an Agilent 7890B gas chromatograph coupled to an Agilent 5977A mass spectrometer. The column was a Rxi-5SilMS from Restek (30m with 10 m integraguard column - ref 13623-127). Oven temperature ramp was 60 °C for 1 min then 3 °C/min to 240 °C for 10 min. Helium constant pressure was 9.43 psi. Temperatures were the following: injector: 250°C, transfer line: 280°C, source: 230 °C and quadrupole 150 °C. The quadrupole mass spectrometer was switched on after a 3.2 min solvent delay time, scanning from 40-400 u. Data analysis was carried out with the MassHunter Quantitative Analysis software from Agilent. Patchoulol was identified by comparison with an analytical standard (Ref 0569-05-95 HWI group). Absolute quantification of patchoulol was performed using a calibration curve prepared with the analytical standard.

## Accession numbers

All genes information used in this article can be found in Phytozome data library (https://phytozome-next.jgi.doe.gov/info/CreinhardtiiCC_4532_v6_1) under the following ID for PRK and RbcS2 respectively: Cre12.g554800_4532 (PAC:52538461) and Cre02.g120150_4532 (PAC:52513304).

## Acknowledgments

This work was financed by the ANR project CalvinReboot (ANR-23-CE44-0007) and supported by grant DIM BioconvS from Region Ile-de-France and by grant Sesame Filiere BFB financed by Region Ile de France and the government as part of the France 2030 Plan. Finally, the authors wish to thank Dr. Tanguy Chotel and Mr. Bruno Da Silva-Teixeira for their valuable insights.

## Author contributions

MB, BG, SDL and PC designed the research; MB, LJ and FG performed the experiments; MB, SDL and PC wrote the manuscript.

## Data availability

All data are presented in the manuscript. Raw data are stored on the Sorbonne Université cloud and are available from the corresponding author upon request. The level 0 intron parts described in this study will be deposited at the Chlamydomonas Resource Center for distribution within the existing *C. reinhardtii* MoClo kit, and their characteristics and sequences are provided in Data S1 and S2 respectively.

## Supporting information

Data S1 contains all the information about the plasmids used in this article.

Data S2 contains all the plasmid maps used in this article.

Figure S1 contains the agarose gel demonstrating correct splicing of the introns PRKi1, PRKi2 and RBCS2i1.

