## Supplementary material for "PRK introns provide modular tools for enhanced transgene expression in *Chlamydomonas reinhardtii*": Figure S1

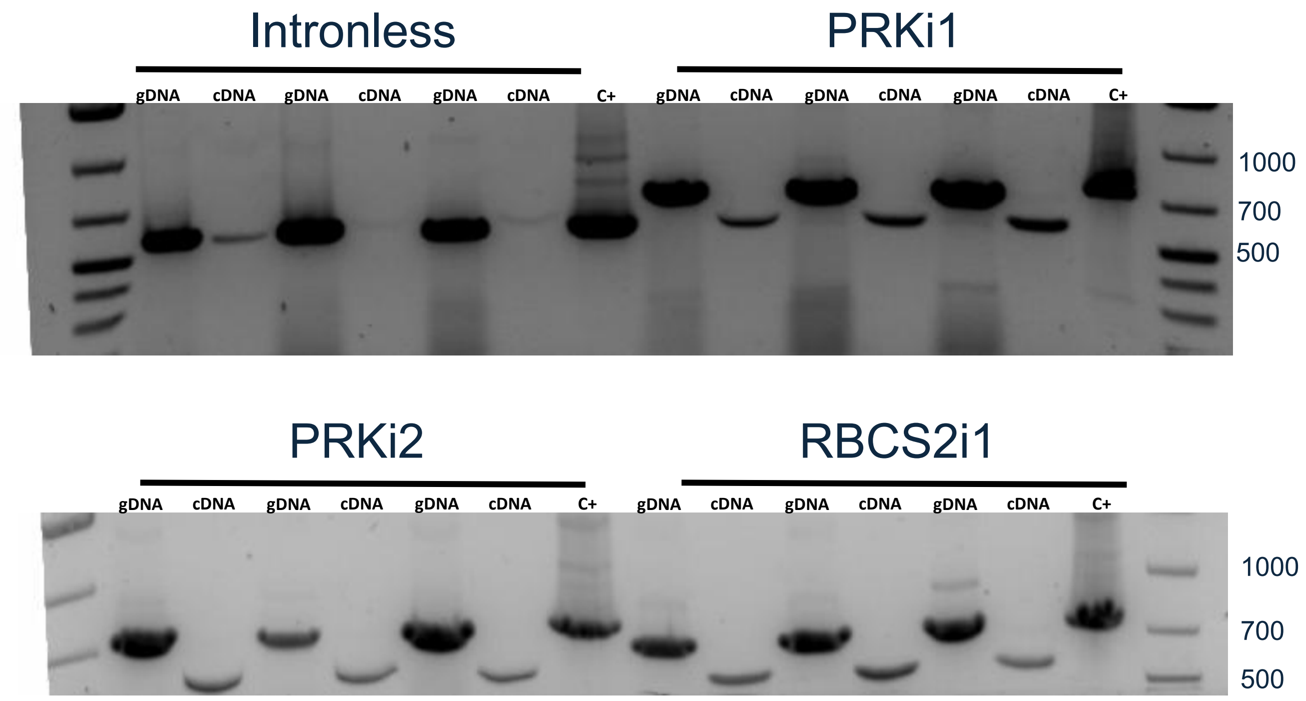


**Figure S1.** Testing of splicing three colonies for each construct containing either no intron, PRKi1, PRKi2 or RBCS2i1. Electrophoresis of PCR products obtained from genomic DNA extracts, cDNA from transformants and plasmid (C+). The expected size for all the amplicons on the cDNA is 562 nt. For the genomic DNA and the plasmid the expected size of the amplicons for Intronless, PRKi1, PRKi2 and RBCS2i1 is 562 nt, 734 nt, 780 nt and 739 nt.
